# How clean is clean? In-vitro comparison of biofilm removal efficacy and cleaning characteristics of three debridement pads

**DOI:** 10.1101/2023.04.19.537160

**Authors:** B. Liepins, C. Tauscher, C. Panhölzl, T. Leeb, C. Häusler, C. Rohrer

## Abstract

**Aim:** The aim of this study was to elucidate the effectiveness of soft debridement in cleaning wounds varying in size and type of exudate and in creating/maintaining a window of opportunity for the wound to be able to heal.

**Methods:** This study presents a standardised in-vitro comparison of three different debridement pads based on the defined composition of exudate and standardised cleaning protocol followed by an robotic cleaning arm. Three important cleaning characteristics (fluid holding capacity, cleaning efficacy/capacity) and the biofilm removal efficacy of wounds varying in size and composition and viscosity of exudate were assessed.

**Results:** All three debridement pads tested showed the ability to clean small to large wounds with different types of exudate (serous/fibrinous) as well as to remove biofilm cells to some extent. Long and dense fibres are favourable when it comes to taking up and holding onto exudate while shorter fibres help to break open harder to clean wounds.

**Conclusion:** A balance between fluid holding capacity and cleaning efficacy/capacity is important in order to achieve the best overall results and successfully remove exudate as well as biofilm cells from small to large wounds with different types of exudate. This in turn has a potential influence on the microenvironment of the wound.

**Key points:**

- The right balance between the parameters tested in this study is crucial for a successful biofilm removal.
- The type of exudate (serous, fibrinous) has an influence on the cleaning efficacy/capacity of debridement pads.
- Soft debridement is able to remove biofilm cells and devitalized tissue as well as dead cells, exudate, proteins etc.
- Good cleaning efficacies without the ability to take up and hold onto exudate, protein and cells are not sufficient for the successful removal of biofilm.

## 1 Introduction

Wounds that are halted in the inflammatory phase of wound healing are called non-healing or chronic wounds. Characteristics of these wounds are the presence of pus and slough, necrotic tissue and bacteria.^1^ The list of reasons for impaired wound healing and chronic wounds is long and reaches from underlying conditions of the patient to inappropriate treatment.^2^ Sharp debridement is known to change the microenvironment of the wound and to create a window of opportunity^3^ in which the wound is more susceptible to treatment and possible healing.

The presence of bacterial biofilms is thought to be frequently linked to non-healing chronic wounds, even though the underlying mechanism is not yet fully understood.^4^ Studies have shown that biofilm communities are present in over 78% of chronic wounds.^5^ Biofilms are aggregates of bacteria often embedded within the wound with a change microenvironment as a result. The biofilm phenotype and the changed microenvironment in the wound results in bacterial tolerance towards antibiotics and antimicrobial agents as well as protection from the immune system. ^6^ Most of our knowledge regarding biofilm are derived form in vitro experiments which are difficult to extrapolate to the in vivo situation however a change in microenvironment and growth induction seem to enable better eradication. ^7–10^

One strategy to support the immune system in combating the biofilm related infection and to increase the success of wound healing treatment is wound debridement. The term “debridement” stands for the removal of bioburden from wounds, including necrotic material, eschar, devitalized tissue, infected tissue, hyperkeratosis, slough, pus, haematomas, debris, bone fragments and foreign bodies. Additionally, it nurtures the wound edges and peri-wound skin where biofilms are most active and where cells that promote epithelialization can be found.^11–13^ Debridement has shown to play an important role in the treatment of wounds infected with biofilm^14,15^ as it removes devitalized tissue, adherent biofilm, surrounding EPS and a multitude of pro-inflammatory substrates. It is the initial step to ensure an optimal wound healing environment by improving the microcirculation in the wound and reducing inflammation and an inseparable element of wound bed preparation.^12^ Most importantly, debridement creates a time-dependent “window of opportunity”, during which biofilms are more susceptible to treatment with antiseptics.^3,8,15^

Types of debridement include enzymatic debridement, ultrasonic debridement (thermal and biological effect), sharp debridement (removal of all devitalized tissue), autolytic debridement (hydrogels, hydrocolloids), and soft debridement (debridement using physical strength).^16^ The most appropriate method should be carefully selected and depends on factors such as tissue type, presence of biofilm, depth and location of the wound as well as the skills of the performing person and the preference of the patient ^12,17,18^. Indications where debridement can be of benefit include acute wounds, diabetic foot ulcers, vascular leg ulcers, pressure ulcers and chronic wounds.^17,18^

Exudation is the process of liquid excretion from the capillary and lymph system in the course of an inflammation process. The type of exudate secreted by wounds tells a lot about the condition and state of the wound. Types of clinical exudate include serous, fibrinous, haemorrhagic and purulent wound fluid. The viscosity of the wound fluid is dependent on the fibrin amount whereas the colouring stands in correlation with the cell content. In early stages serous wound fluid is excreted which is usually amber, runny and clear. Infected wounds excrete fibrinous wound fluid, which is usually milky, creamy and viscous.^19^

Over the last couple of years several soft debridement tools (pads, foams, cloths) have been introduced into the market. They differ in material composition but are used in a similar manner. Debridement pads differ in type, number, length and cut of the fibre used on the wound facing side while sponges differ in density, strength and pore size. These properties all influence the possible exudate uptake as well as the cleaning success.^20^ This study focuses on the influence of type, length and density of fibres on the cleaning success, therefore only debridement pads were reviewed. The debridement pads should be prewetted with fluid (e.g. normal saline, sterile water, wound cleansing solution) before providing a circular motion with gentle, ongoing pressure. Premoistening of the pads enhances debridement and decreases pain.^16^

The aim of this study was the characterization of three mechanical debridement pads regarding their performance in fluid holding capacity, cleaning efficacy and cleaning capacity and finally their performance in biofilm removal of four different organisms.

## 2 Materials and Methods

### 2.1 Materials

Gelatine fix powder (Dr.Oetker, Bielefeld, Germany) and collagen powder (Nature Diet, Premium food) were purchased at Amazon. Egg white powder from chicken (EO500, MERCK, Darmstadt, Germany) was purchased from Carl Roth. 75% Cochenille red (C.I: 16255, solid, PanReac AppliChem, Darmstadt, Germany) was purchased at Carl Roth. ASTM Typ 2 water was used for all experiments. The acrylic base plates (acrylic sheet, 8×8cm) were purchased at Perspex. Test organisms *Escherichia coli* ATCC 8739, *Enterococcus faecalis* ATCC 29212, *Staphylococcus epidermidis* ATCC 12228 and *Staphylococcus aureus* ATCC 6538 were purchased as Culti-Loops^TM^ at Thermo Fisher Scientific. Tryptic soy broth (TSB, Merck) was purchased at Avantor and ready to use tryptic soy agar (TSA) plates were purchased at bioMérieux. Sterile 0.2 μm cellulose nitrate membrane filters with a diameter of 47mm were purchased at either Whatman® or Sartorius. Sodium chloride was purchased at Carl Roth and centrifuge Tubes, tissue culture cell scrapers and 1.5ml Eppendorf tubes were purchased via Avantor. Three different mechanical debridement pads currently on the market were assessed. Debrisoft® pad (DP, Lohmann and Rauscher GmbH&Co.KG) is a sterile, single-use monofilament fibre pad made of a densely packed and angled polyester fibres. The rear side of the product has a grip pocket made of polyester, polyamide and elastane. The pad needs to be soaked with 20-40 ml of water or saline before use (acc. to IFU). Cutimed® DebriClean (CDC, BSN medical GmbH) is a sterile, single use debridement pad, whose wound side consists of white mono-filament microfibre loops and blue, abrasive microfibre loops. According to the IFU it should be fully saturated with wound cleansing solution (appr. 30 ml) before use. Prontosan® Debridement pad (PDP, B. Braun Medical AG) is a sterile, single use debridement pad consisting of microfibres that debride and an absorbent backing layer made of polypropylene and polyester. According to the marketing materials the unprinted side should be moistened with 15-20 ml of Prontosan® wound irrigation solution before use.

### 2.2 Serous/fibrinous wound fluid simulation

Two types of wound fluid simulating serous and fibrinous wound exudate were used throughout the course of this study. The serous wound fluid simulation contains egg white powder from chicken (10% (w/w)) in water, which is heated to 40°C under constant stirring until the powder is completely dissolved. To enable visual analysis throughout the experiments Cochenille red (0,05 % [w/w]) is added to the mixture. The fibrinous wound fluid simulation is prepared by dissolving gelatine (18,9 % [w/w]) and collagen (37,7 % [w/w]) in H_2_O in two separate glass vessels. The solutions are heated to 40°C and constantly stirred. Once completely dissolved, the mixtures are combined and Cochenille red (0,05 % [w/w]) added. The fibrinous wound fluid simulation needs to be pipetted while still warm as the viscosity increases once cooled down to room temperature.

### 2.3 Fluid holding capacity (FHC)

The fluid holding capacity was assessed with both types of wound fluid according to ^21^. The debridement pads were weighed (mDRY1 [g]) before dipping them into 120ml of wound exudate simulation for 60 seconds with the fibre side of the product. The soaked product is immediately weighed lying flat in a large petri dish (mWET1 [g]) before allowing excess fluid to drip off by holding it upright with a pair of tweezers for 30 seconds. After weighing the product (mWET2 [g]) leave to dry in a drying oven (70°C, ventilation open) for at least 4 hours. The dry debridement pad is weighed again (mDRY2 [g]).

FHC1 (1) shows how much wound fluid/exudate including protein, cell debris etc. the pad is able to hold.

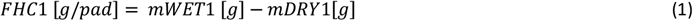

FHC2 (2) describes how much wound fluid/exudate including protein, cell debris etc. the pad is able to hold on to after being held vertically for 30 seconds.

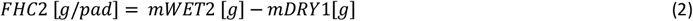

The total capacity gives insight on how much protein/cell debris the respective debridement pads can absorb.

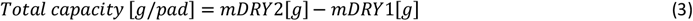

### 2.4 Preparation of test plates

The surface of the 8 x 8 cm acrylic glass plates was treated with a laser cutter (Universal Laser Systems, Versa Laser VLS3.50) to create a round indentation with a diameter of 6 cm with a rough surface. Each plate is marked with a unique number to ensure traceability. 4 ml of the respective wound fluid simulation where pipetted into the round indentation created by the laser cutter, which simulates the wound surface. A uniform distribution of the fluid is achieved by spreading out the liquid using a glass rod. The test plates containing either serous or fibrinous wound fluid simulation are left to dry at constant room temperature and humidity for 2h/24h, respectively. Each plate is weighed before applying the wound fluid simulation (p1 [g]) and after the respective drying time (p2[g]).

### 2.5 “DebriSim”-wound debridement model

The “DebriSim”, a robotic debridement simulation device as described in ^22^ was used for this study. The robotic arm is able to simulate debridement processes by mounting debridement pads to it and cleaning glass plates with different types of wound fluid on it. The type of movement (pattern), pressure applied (weight), number of cycles and speed of movement can be controlled.

### 2.6 Cleaning efficacy

The cleaning efficacy was assessed by weighing the dry debridement pads (d1 [g]) before prewetting them with 20 ml (DP), 30 ml (CDC) and 15 ml (PDP) of H_2_O. The respective amount of liquid was pipetted onto the debridement pads allowing excess fluid to drip off, if necessary. The respective debridement pad is mounted on the sample holder of the DebriSim, the test plate inserted and cleaned using the following settings: Cycle = 1, Weight = 2,5 kg, Pattern = 3 (Lissajous 3:4), Speed = 20 mm/s. Both the plate and the debridement pad are removed, weighed again (p3 [g], d2 [g]) and left to dry at room temperature for 1h and 25 hours, respectively. After weighing the dry plate and debridement pad (p4 [g], d3 [g]) the cleaning efficacy and the exudate uptake are calculated using the following equations (4), (5):

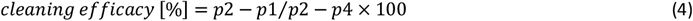

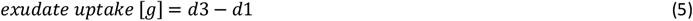

### 2.7 Cleaning capacity

Preparation of plates and debridement pads according to 2.6.

When assessing the cleaning capacity, the same debridement pad is used to clean 5 test plates in a row. The test plates and the debridement pad are weighed after the experiment (p3 [g], d2 [g]) and left to dry according to 4.4. After weighing the dry plate and debridement pad (p4 [g], d3 [g]) the cleaning capacity and exudate uptake are calculated (4,5).

### 2.8 Biofilm removal

Stock cultures (*E. coli* ATCC 8739, *E. faecalis* ATCC 29212, *S. epidermidis* ATCC 12228, *S. aureus* ATCC 6538) were started by inoculation of a Culti-Loop® in TSB and incubation at 35 °C ± 2 °C for 16 – 24 h. An overnight culture was prepared by transferring 100 μl of stock culture into 7ml TSB and incubation at 35 °C ± 2 °C for 16 – 24 h. From the overnight culture a 1:500 dilution in TSB was prepared and incubated at 35 °C ± 2 °C for 1h in a horizontal shaker. Sterile 0.2 μm cellulose nitrate membrane filters with 47 mm in diameter were put onto TSA plates. Each membrane filter was spiked with 9 x 2 μl of the diluted working culture. The TSA plates containing spiked membrane filters were incubated at 35 °C ± 2 °C for 72 h. During the incubation period every 16 – 24 h membrane filters with established biofilm were transferred onto fresh TSA plates. After 72 h membrane filters with established biofilm were transferred onto regular glass plates (28×28 cm) that were treated with a laser cutter to create a round indentation with a diameter of > 6 mm with a rough surface. To increase adhesion of membrane filters to the glass surface each plate was covered with a thin film of gelatine solution (25% w/v) and left to dry for 30 – 60 min. At first 100 μl of sterile water were pipetted onto each glass plate and then a membrane filter with biofilm was put onto the wetted gelatine surface. Next the membrane filters were covered with 4ml of fibrinous wound fluid simulation at just below 40 °C and left to dry for 90 – 120 min. Biofilm glass plates were placed into the DebriSim and a pre-wetted debridement pad (20 ml (DP), 30 ml (CDC) and 15 ml (PDP)) was mounted into the sample holder. Debridement was simulated using the following settings: (Cycles = 6, Weight= 0.25 kg, Pattern = 3, Speed = 20 mm/s). A non-debrided control was included and measured for each strain. After debridement each membrane filter was removed from the glass surface and transferred into 50 ml centrifuge tubes (Avantor) containing 36 ml sterile 0.9 % (w/v) sodium chloride solution. Remaining fibrinous wound fluid simulation on the glass surface was collected with cell scrapers for tissue culture (Avantor) and added to the membrane filter. Centrifuge tubes were treated in an ultrasonic bath (USC 1200T - Avantor) for 5 min and then vortexed (Analog Vortex Mixer - Avantor) for 30 sec. Dilutions of each sample were prepared in 1.5ml Eppendorf reaction tubes and 100μl of each dilution were spread onto TSA plates. The plates were incubated at 25 °C ± 3 °C for 72 – 96 h. Lastly all forming colonies were counted and the results prepared for statistical analysis.

### 2.9 Statistical analysis

All statistical analysis were performed in R ^23^ Statistical Software (version 4.0.4; R Core Team 2021). Tables containing raw data were imported with the *readxl* ^24^ package. Data was processed and prepared for analysis using the *tidyverse* ^25^and *magrittr* ^26^packages. A Shapiro-Wilk test was done using *rstatix* ^27^and Levene’s test was performed using *car* ^28^. A one sided ANOVA and Tukey’s HSD were done with the *stats* ^23^package to elucidate significant differences between groups. Adjusted p-values were plotted into the graphs with help of *ggpubr* ^29^package. P_adj-values if shown were labelled as follows: p <= 0.001 – ‘***’, p <= 0.01 – ‘**’, p <= 0.05 – ‘*’, p > 0.05 – ‘ns’;

## 3 Results

### 3.1 Fluid holding capacity (FHC)

The FHC, as the name implies, gives insight on how much exudate a debridement pad is able to absorb and is crucial for the overall debridement success. Transferred to the clinical practice it influences the cleaning success of the wound and the number of pads needed to achieve a “clean” wound. As the debridement pads can only be used as a whole, all results are presented per pad instead of per g material. For reference, DP is the heaviest with an average of 10.6 g, while CDC and PDP weigh 3,8 g and 2,5 g, respectively. The results for FHC1 and FHC2 shown in Figure 2 follow the same trend. Independent of the composition of the wound fluid, DP is able to absorb the most fluid. It is followed by CDC and PDP. The composition and viscosity of the wound fluid does influence the calculated parameters, as they are between 10 and 25% higher when fibrinous wound fluid simulation is used. The overall higher FHC values for the fibrinous wound fluid simulation can be explained by the higher density of the liquid, resulting in a higher weight taken up per ml of wound fluid. DP exhibits the highest data consistency throughout the experiments. It shows the lowest difference (10%) in FHC1 and FHC2 when comparing the two types of wound fluid simulations, which indicates that the majority of exudate taken up by DP stays in the pad.

**Figure 1:**
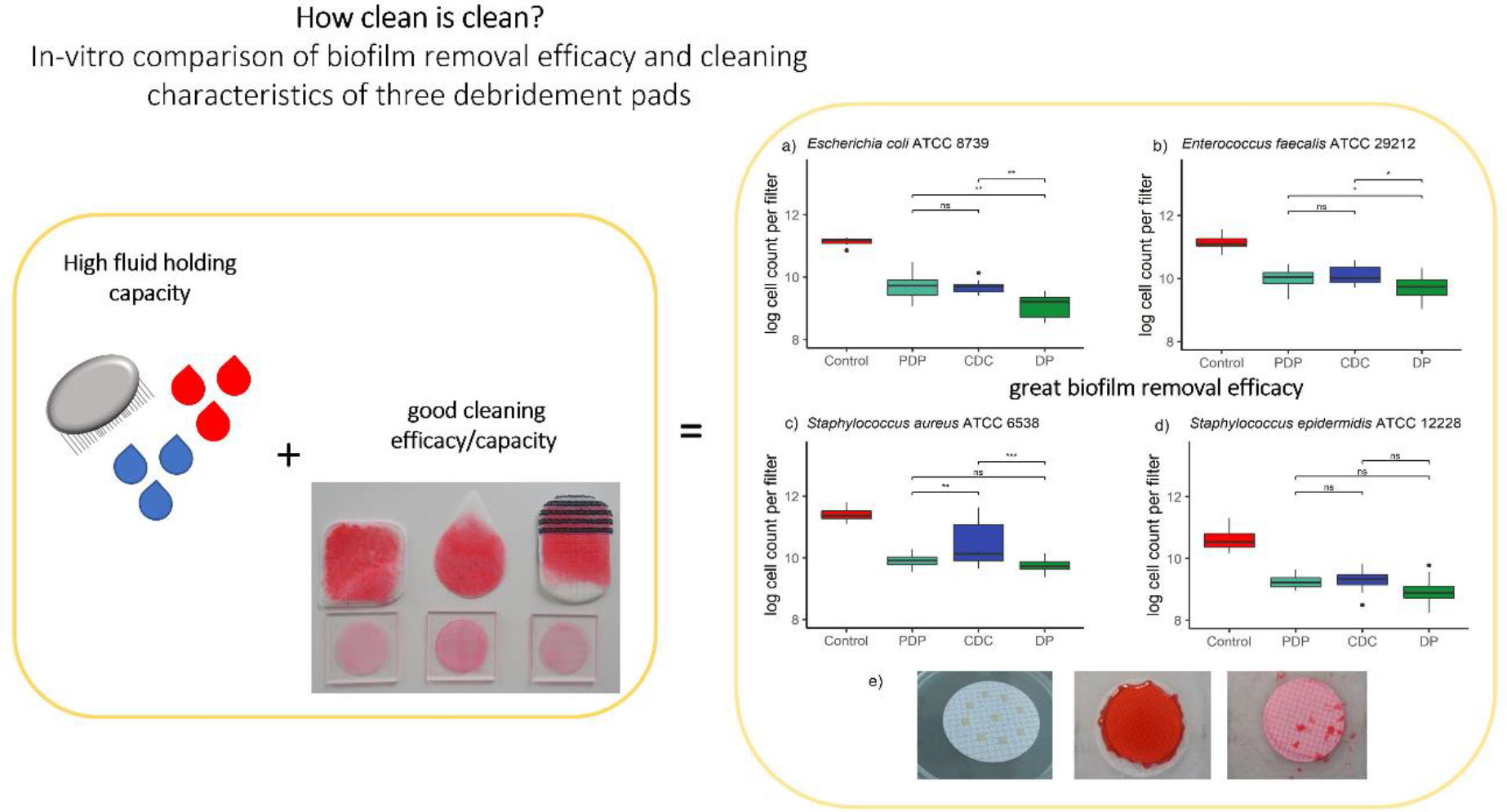
Graphical abstract.

**Figure 2.**
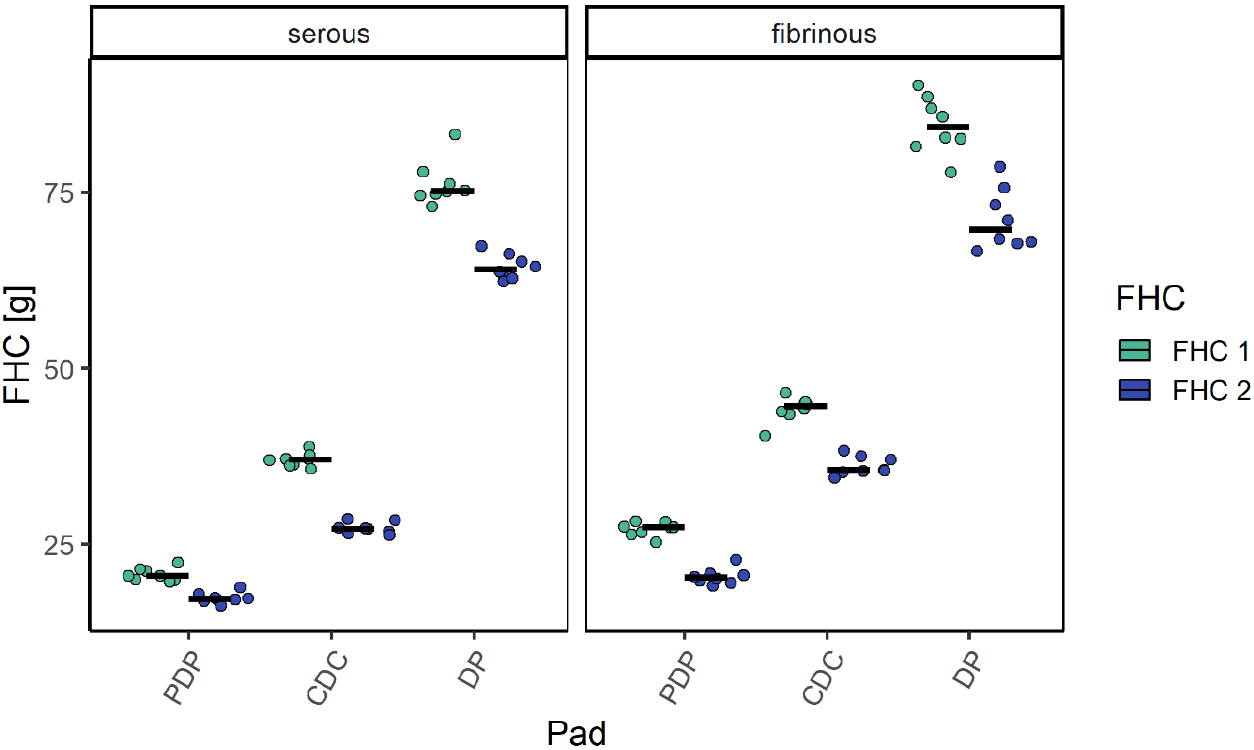
displays the results of FHC1 [g] and FHC2 [g] for all three debridement pads (PDP, CDC and DP) tested with serous and fibrinous wound fluid simulation in a scatter plot. The green and blue points show the single values while the black line represents the median.

The trend observed in the total capacity is the same as in the FHC1/FHC1 experiments. DP absorbs the most protein, followed by CDC and PDP. As to be expected the total capacity was much higher (around 70%) when the fibrinous wound fluid was tested compared to the serous wound fluid simulation. The difference can be explained with the protein content in the fibrinous wound fluid simulation being nearly 60% as compared to only 10% the serous wound fluid simulation.

### 3.2 Cleaning efficacy

All three pads perform equally well when testing the cleaning efficacy [%] with serous wound fluid simulation (Figure 4). There is no significant difference between the pads as they were all able to remove around 92% of the wound fluid from the respective plates. Testing the cleaning efficacy with fibrinous wound fluid simulation showed, that using the CDC only resulted in an average of 62% cleaning efficacy. PDP was able to remove an average of 92% of wound fluid simulation outperforming DP by nearly 11%, while both had a significant difference to CDC. The results for PDP were the most consistent followed by DP and CDC.

**Figure 3.**
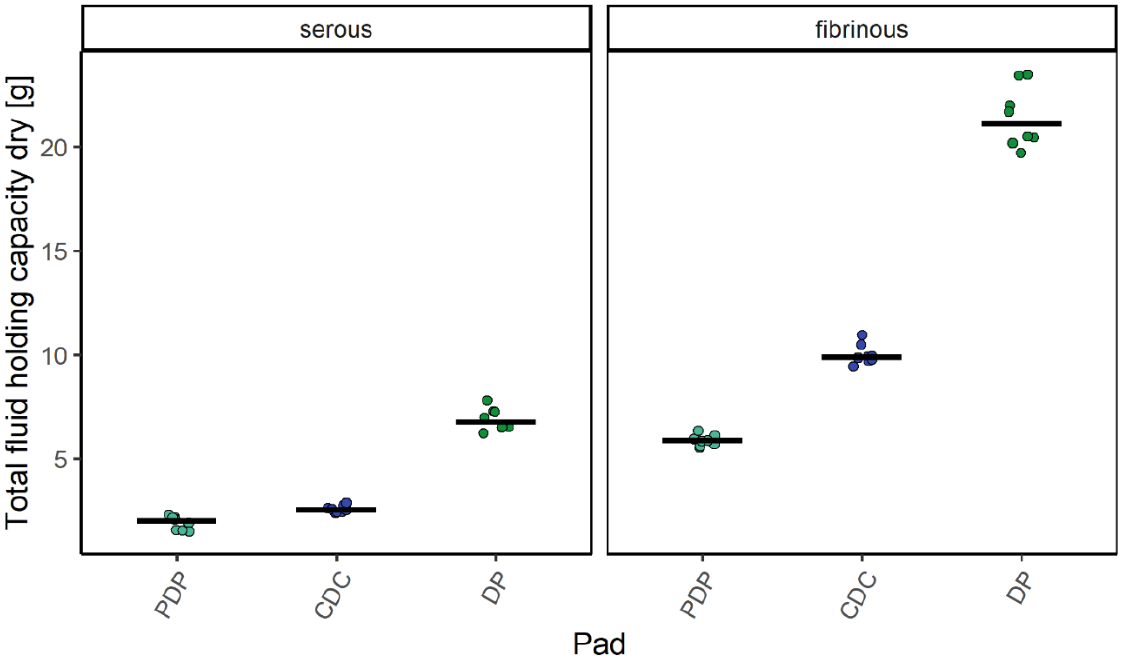
shows a comparison of the results for the total capacity [g/pad] of all three debridement pads (PDP, CDC and DP) tested with serous and fibrinous wound fluid simulation in a scatter plot. The green and blue points show the single values while the black line represents the median.

**Figure 4.**
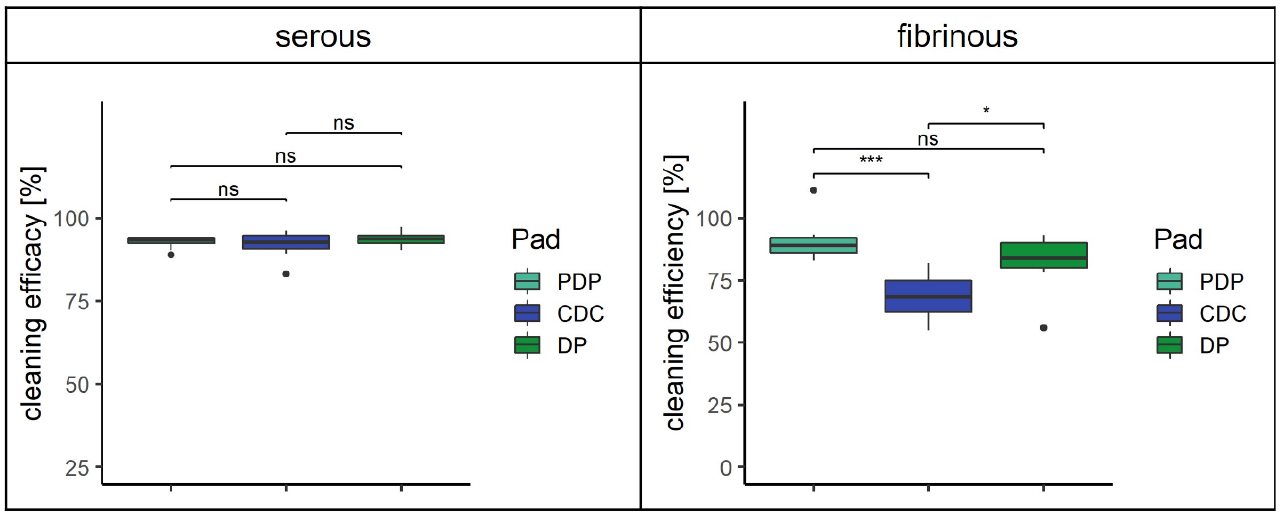
shows a comparison of the results for the cleaning efficacy [%] of all three debridement pads (PDP, CDC and DP) tested with serous and fibrinous wound fluid simulation in a box plot. P_adj-values were calculated for all groups and labelled as follows: p <= 0.001 – ‘***’, p <= 0.01 – ‘**’, p <= 0.05 – ‘*’, p > 0.05 – ‘ns’.

### 3.3 Cleaning capacity

For the cleaning capacity five plates are cleaned in a row without changing the debridement pad mounted to the DebriSim arm. This is to test the ability of the debridement pads to clean bigger wounds and/or higher wound fluid levels compared to the cleaning efficacy.

The results of the cleaning capacity for serous wound fluid (Figure 5) compared to the results of the cleaning efficacy (Figure 4). All three pads performed well with the best and worst result lying only 6% apart. The only significant difference presented itself between DP and PDP with p_adj <= 0.05.

**Figure 5.**
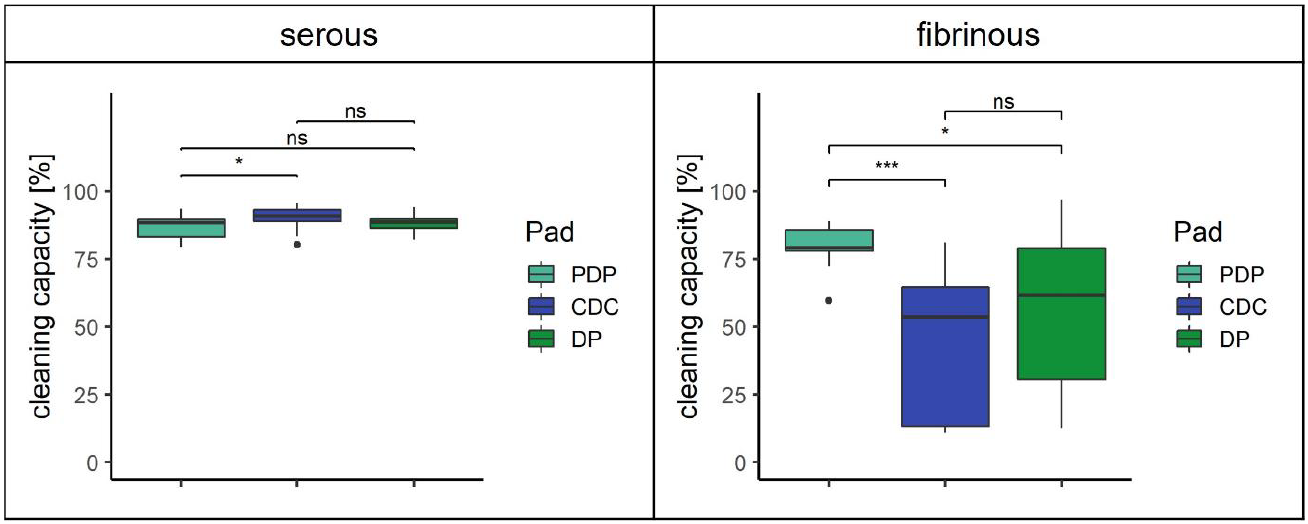
shows a comparison of the results for the cleaning capacity [%] of all three debridement pads (PDP, CDC and DP) tested with serous and fibrinous wound fluid simulation in a box plot. P_adj-values were calculated for all groups and labelled as follows: p <= 0.001 – ‘***’, p <= 0.01 – ‘**’, p <= 0.05 – ‘*’, p > 0.05 – ‘ns’.

The cleaning capacity for fibrinous wound fluid simulation equals a large, hard to clean wound with firm slough. PDP achieved an average cleaning capacity of 75% with a narrow distribution of results. The difference to DP and CDC were both significant. The results also show that the variance of the results increases with wound size and viscosity and stickiness of the wound fluid. With increasing saturation of the pad, the ability to clean decreases but due to the pressure and distinct motion of the DebriSim arm, bigger parts of the wound exudate, especially in the case of fibrinous wound fluid, break off and are pushed off the test plate. This doesn't occur in a reproducible manner, which leads to big deviations in results rather quickly.

When looking at the mean exudate uptake [g] per pad (Figure 6), the results don't always correlate with the cleaning efficacy and capacity. Product uptake when testing the efficacy with serous wound fluid was nearly equal for all pads with very reproducible results. Results started to spread when testing the product uptake with fibrinous wound fluid simulation but the average results stay close together. It shows very well, that the saturation with exudate is not reached after cleaning a small wound (=cleaning efficacy). DP was able to take up the most exudate in both cases which confirms the results from the FHC experiments. Interestingly, even though PDP was able to clean off fibrinous wound fluid from larger wounds, it takes up the least exudate out of the three pads. This leads to the conclusion that the majority of exudate is pushed to the surrounding area of the test plate.

**Figure 6.**
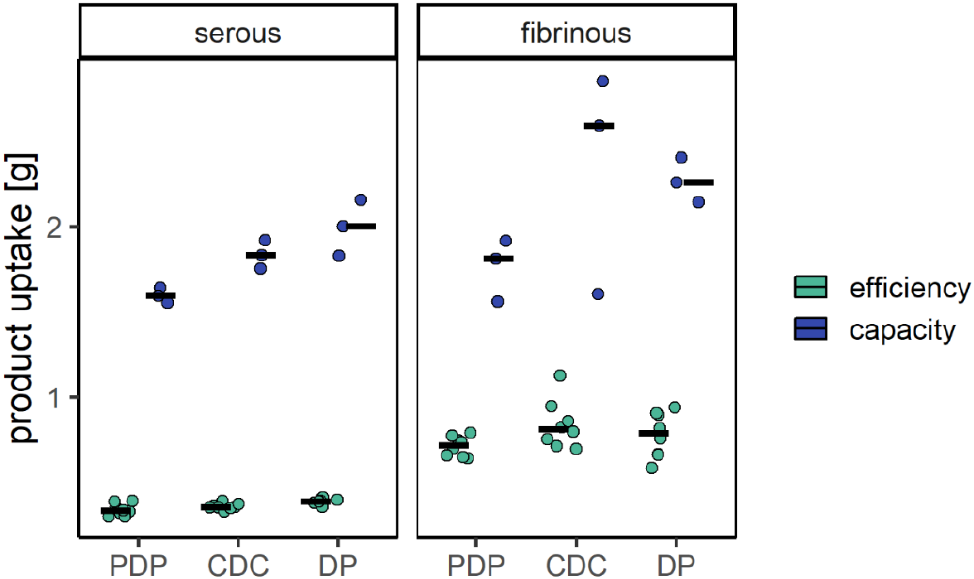
shows the results for the mean exudate uptake [g] of all three debridement pads (PDP, CDC and DP) tested with serous and fibrinous wound fluid simulation in a scatter plot. The green and blue points show the single values while the black line represents the median.

### 3.4 Biofilm removal

In this experimental setup the established biofilm was covered with fibrinous wound fluid simulation in order to create “wound like” conditions. This means that the fibrinous wound fluid simulation has to be removed before the biofilm can be reached. The same settings as for cleaning efficacy and capacity were used on the DebriSim to achieve comparable results. According to the IFU DP can be prewetted with a volume ranging from 20-40 ml. Preliminary tests showed (data not shown), that the prewetting volume does not influence the outcome of the previously tested parameters. To compare how well the debridement pads are able to remove biofilm based on their structure and composition, water was used to prewet them.

Biofilms by both gram negative and gram positive bacteria were tested as shown in Figure 7 a-d. Overall DP was able to remove the most biofilm cells independent of the organism, showing the most significant differences to PDP and CDC. PDP was able to remove *S. aureus* biofilm significantly better than CDC, whereas in all other organisms no significant difference could be achieved between these two debridement pads. DP achieved the best results in *E. coli* and *S. aureus*, with the most significant differences being measured in *S. aureus. S. epidermidis* was the only species, where no significant difference between the three pads could be measured.

**Figure 7.**
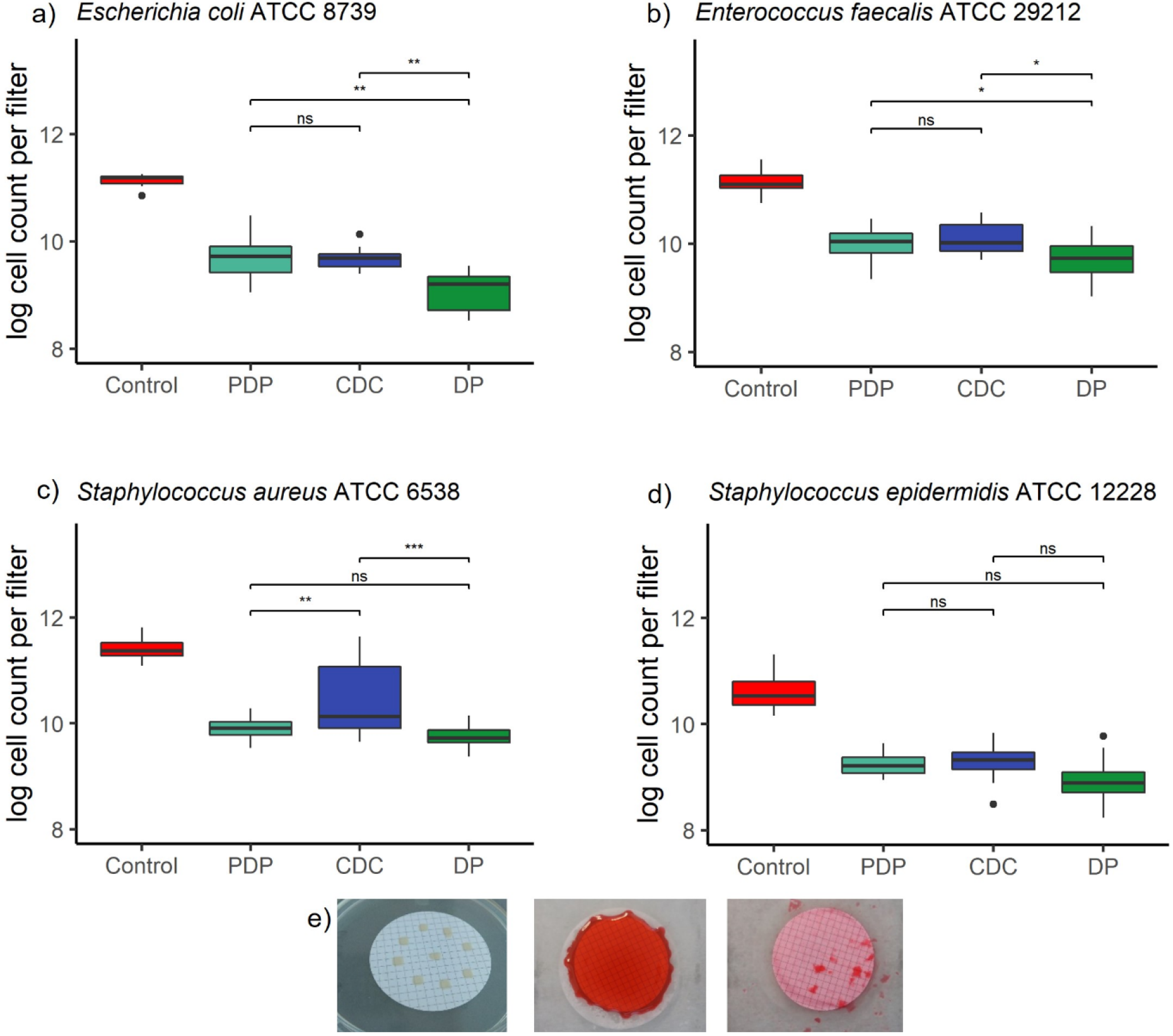
shows a box plot of the log cell count per filter compared to the control for PDP, CDC, DP testing the removal of mature biofilms from for the following organisms: E.coli (a), E. faecalis (b), S. aureus (c) and S. epidermidis (d). P_adj-values were calculated for all groups and labelled as follows: p <= 0.001 – ‘***’, p <= 0.01 – ‘**’, p <= 0.05 – ‘*’, p > 0.05 – ‘ns’. Part e) shows exemplary photos taken of a filter with mature biofilm in dots, the biofilm covered with stained fibrinous wound fluid simulation and the cleaned filter after the debridement process (from left to right).

## 4 Discussion

Chronic wounds are notoriously difficult to treat, but debridement has shown to be one of the beneficial tools for improved healing. Debridement pads for soft debridement have been developed as an easy debridement tool. Several features of the debridement pads determine its efficiency and efficacy. The FHC is one of the most important features but differs between the tested pads. The measured differences in FHC, in this study are due to the composition and setup of the debridement pads. The most obvious difference between DP and CDC (PDP) is the length and density of the fibres present. The setup of DP with its densely packed and long, fibres is able to deal with different types of fluids in a very consistent manner, absorbs the most fluid and retain it. The short and thin microfibres on the PDP, only absorb around one third of the exudate compared to DP. The in-vitro testing of the cleaning efficacy translates to the cleaning of a small wound in the clinical practice and shows how well the debridement pads remove the respective wound simulation from the test plate. The two types of wound fluid used in this study represent the two ends of the wound fluid spectrum. The serous wound fluid simulation represents easy to clean wounds with low viscosity exudate and the fibrinous wound fluid simulation a hard to clean wound with very viscous exudate. The more viscous the exudate the harder the wound is to clean. As the wound fluid simulations are left to cool on the plates before the cleaning rounds, the viscosity increases, and the surface becomes dry to the touch, forming a sort of protective cover. The “wound exudate” of the fibrinous wound fluid simulation has a higher viscosity and stickiness and is more difficult to break apart. The cleaning pattern was repeated 5 times by the DebriSim arm within one cycle. Throughout the process the wound fluid simulation was usually not evenly removed round by round as the first 1-3 rounds are needed to break the surface open, before it can be removed from the plate. The removed wound fluid simulation is not completely retained by the debridement pads but also pushed to the surrounding surfaces. Overall, it was shown that all three pads tested are suitable for the debridement of small and large wounds presenting various types and amounts of exudate, but that consistency of results strongly varies and that cleaning efficacy and capacity don't directly translate into retention. The long and dense fibres of DP are able to break open the biofilm matrix as well as take up the cells within the biofilm and remove it. The results show that DP is able to remove biofilms containing both gram negative and gram positive bacteria, which is crucial as wound biofilm always contain several species. The removal efficacy for the different organisms varies as the composition and also properties of the biofilm itself changes depending on the species. As all of the pads were able to mechanically remove the biofilm cells of *S. epidermidis* equally well it suggests that this organism forms an in vitro biofilm with low adhesion to the membrane and/or a weak and easy to remove EPS matrix structure. It is crucial for a soft debridement pad to combine high FHC with sufficient cleaning efficacies as DP shows the best overall results in the biofilm removal experiments.

In conclusion, we here show that soft debridement can clean wounds ranging from easy to hard to clean as well as remove biofilm cells and therefore could have a potential influence on microenvironment of the wound. Soft debridement can maintain a healthy wound after sharp debridement and might even keep a wound from getting into a state where sharp debridement becomes necessary.

### Disclaimer

This work has been supported by ACMIT - Austrian Center for Medical Innovation and Technology, which is funded within the scope of the COMET - Competence Centers for Excellent Technologies program and funded by the federal government (BMWA und BMK) and the governments of Lower Austria and Tyrol.

## Abbreviations and acronyms

FHC: fluid holding capacity
DP: Debrisoft® Pad
PDP: Prontosil® Debridement Pad
CDC: Cutimed® DebriClean
EPS: Extracellular polymeric substances
TSB: Tryptic soy broth
TSA: Tryptic soy agar
IFU: Instruction for use
*S. aureus*: Staphylococcus aureus ATCC 6538
*S. epidermidis*: Staphylococcus epidermidis ATCC 12228
*E. coli*: Escherichia coli ATCC 8739
*E. faecalis*: Enterococcus faecalis ATCC 29212

## Notes

### Competing Interest Statement

The authors have declared no competing interest.

### Summary of Updates

Author Cornelia Haeusler, a disclaimer and graphical Abstract was added. The structure of the abstract and overall publication was slightly changed.

